# Nest attributes influence choice accuracy, but not decision speed in acorn ants

**DOI:** 10.1101/2025.07.20.665794

**Authors:** Sheila Shu-Laam Chan, Claire Hemingway, Isaac P Weinberg, Philip T Starks

**Affiliations:** Department of Biology, Tufts University; Department of Ecology & Evolutionary Biology, University of Tennessee, Knoxville; Collaborative for Animal Behavior, University of Tennessee, Knoxville; Department of Psychology, University of Tennessee, Knoxville

## Abstract

Decision making can have significant fitness consequences across various aspects of animal life. For acorn ants, *Temnothorax curvispinosus*, choosing a new nest quickly and accurately can affect the survival and fitness of the whole colony. When emigrating, ants consider several nest attributes such as cavity shape, thickness, and brightness. Ants may benefit from having more attributes differentiating potential nests only if they can quickly and accurately assess all possible attributes and make well-informed decisions. Here, we asked if the number and type of attributes differentiating potential nests affect the accuracy and latency of colony decision-making. We used pair-wise tests, where potential nests differed in 1-3 attributes, with one nest within the pair considered less optimal. We recorded the nest the colonies chose and the time it took to make the decision. We found that the degree of difference did not affect the decision-making latency, suggesting that ant colonies searching for a new nest might be constrained temporally when selecting a new nest site. We also found that accuracy increased with the number of attributes, particularly when nest brightness was manipulated, indicating that increasing the number of attributes may help facilitate nest-site selection.

## Introduction

Animals routinely make decisions with fitness consequences, such as selecting a mate or deciding what to eat or where to live (1,2). Choices across a variety of contexts are often characterized by multiple attributes. For instance, when selecting a flower to visit, pollinators attend to multiple aspects of nectar rewards, including concentration, volume, variance, and handling time (3,4). In most cases, these attributes are informative about different properties of choices, and attending to them may lead to better decision outcomes (5,6). However, access to more information about potential options will only benefit decision-makers if they can quickly and accurately identify the various attributes of available options and make appropriate decisions using that information.

These complex decision scenarios are particularly common in acorn ants (*Temnothorax sp.*), which live in cavities like fallen acorns or hickory nuts, primarily on the forest floor. Given the relative instability of these nesting sites, acorn ants quickly respond to changing conditions, relocating to a new nest when their current one becomes unsuitable (7,8). During emigration, ants likely have several potential nests to choose from, as multiple acorns have dropped from a single tree at any given time. When confronted with multiple nests, acorn ants send scouts to carefully assess these options by examining several nest attributes, including brightness, cavity shape and size, and entrance width. Previous studies suggest that an ideal nest for acorn ants is dark, with a spacious, thick cavity and a narrow entrance (9,10). This complex nest-selection process requires ants to integrate information within and across potential nest sites. Scouts then recruit more ants to preferred nest sites via tandem running (11). When a quorum of ants considers a nest acceptable, recruitment shifts from tandem running to direct transport of the remaining colony members, including the brood and queen(s). As the number of ants in the new nest increases, the rate of the emigration process also increases (i.e., the ants collectively decide on a new nest via positive feedback) (8). During this process, the quality of the candidate nest affects how quickly the ants decide to initiate recruitment, with better nests typically eliciting faster recruitment and migration (7,12).

Ants may benefit from having more distinguishing attributes among potential nests only if they can efficiently identify available nest sites, evaluate the attributes of each, and make well-informed decisions based on this information. Increasing variation across multiple attributes of nests also has the potential to make decision-making difficult in several ways. First, the more attributes a nest has, the more information ants must process (5,6). For humans, choosing based on a single attribute, like color, is often easier than evaluating options that differ in color, shape, and size (13). Second, not all attributes may be equally important. Previous work has shown that ants evaluate and prioritize attributes differently: brightness appears most important, followed by the shape of the cavity and, finally, the entrance width (9). Ants also learn to rely on more informative attributes that reliably predict nest quality (14). For instance, in an experimental setting, when only one attribute differentiated options, ant colonies increased their reliance on this attribute, relative to others (14). Third, decisions become more complex when attributes conflict and no single option is considered the best (15,16). Here, choosing a nest based on a good value for one attribute may result in a bad value for another.

When multiple attributes characterize nests, this may have important consequences for decision accuracy and speed. Accuracy may improve when additional attributes increase a nest’s relative value (17). However, in complex choice environments, individuals often rely on decision shortcuts, only considering some available information, which may result in biased or context-dependent choices (6,16,18). Individual acorn ants make context-dependent decisions when making multi-attribute decisions with two conflicting nest attributes (19). Similarly, decision speed can decrease as additional attributes make one option easier to choose (17) or increase because processing more information takes longer. Some decision models predict longer decision times as more information is required to reach a threshold (5,6). Most studies of nest site selection focus on choice accuracy, and decision latency may be an equally important outcome to consider when evaluating decision-making outcomes, especially with multi-attribute choices. Thus far, investigations into decision speed have been limited to scenarios where only one attribute - brightness - varied between candidate nests (12,20). Both speed and accuracy are critical components of decision-making, with significant implications for fitness, and animals often face a tradeoff between these two factors (21).

Here, we explored how the number and type of differentiating attributes influence choice accuracy and speed. We gave ant colonies two-choice tests in which potential nest sites differed in either one, two, or three attributes, and we varied the type of attributes that differentiated potential nest sites. We measured which nest the colony chose, how long it took to make the decision, and, in some cases, whether colonies failed to make a choice. We considered several hypotheses about how manipulating attribute number and type may influence choice accuracy and latency. One possibility is that ants will perform better with access to more information about nest attributes, facilitating quick and accurate decisions. Conversely, increasing the information that ants must attend to may increase the time and processing necessary to identify the preferred nest site, leading to longer and less accurate nest selection. Finally, if ants attend primarily to one attribute more than others, we may see similar patterns of decision speed and accuracy, regardless of the number of other attributes distinguishing potential nest sites.

## Methods

### Study system

Acorn ants (*Temnothorax curvispinosus*) are commonly found in eastern North America forests. An average colony of acorn ants consists of around 80 workers, but can range from 10 to 350 individuals (22). As their common name suggests, they are cavity-dwelling ants living in fallen acorns or hickory nuts on the forest floor. We collected 10 *T. curvispinosus* colonies, which ranged in colony size from 14-35 workers. Colonies were collected during the summer of 2021 in Middlesex Fells Reservation in Eastern Massachusetts and maintained in the lab for nine months during experiments. Using a standard protocol, we provided artificial nests (5 cm × 2 cm × 2 mm) with a glass cover, a metal screen base, and wood walls as their home nests (e.g.,(9)). While in captivity, colonies were fed with water, a 1:1 water-sucrose solution, and mealworms, which were replenished once a week.

### Experimental overview

We manipulated three nest attributes in this study: 1) nest brightness (bright or dark), 2) internal height of the cavity (thick or thin), and 3) width of the nest entrance (narrow or wide). Dark nests were covered by a piece of wood to block all light. Bright nests were covered with a transparent microscope slide and exposed to an LED lamp that did not generate heat. The thick and thin nest cavities were 1.5 mm and 3 mm in diameter, respectively. A narrow entrance was 2 mm, and a wide entrance was 4 mm. The dimensions were determined based on previous experiments (S1 Table). All nests were handmade with wood and a glass cover and had approximately similar volumes (Fig 1). The nests had the same dimensions on the outside.

**Figure 1.**
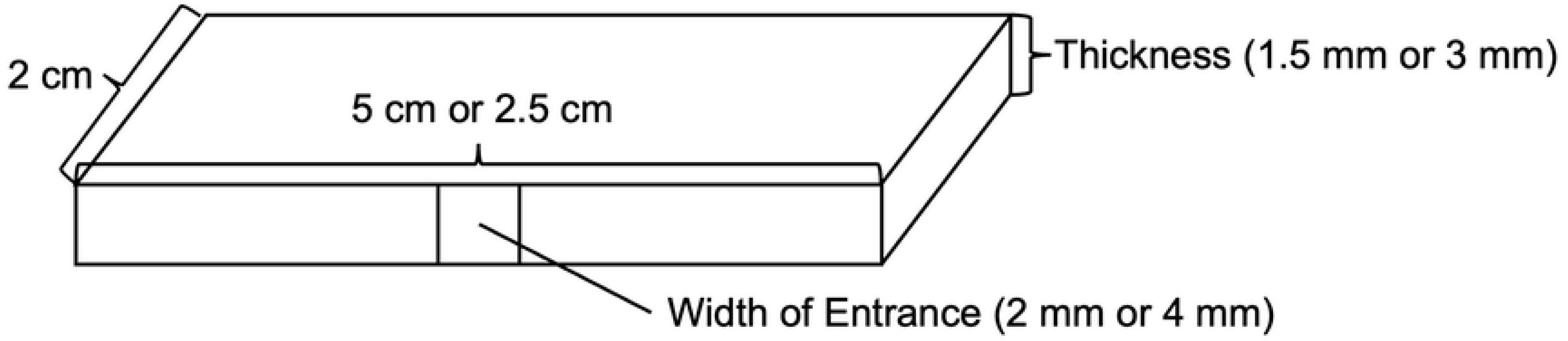
Illustration of nests used in the study. A thick nest was 2.5 cm × 2 cm × 3 mm, and a thin nest was 5 cm × 2 cm × 1.5 mm. The dimensions of thick and thin nests are different to control for the volume of nests. A narrow opening was 2 mm, and a wide opening was 4 mm. A dark nest was covered by a piece of wood and a glass microscope slide, and a bright nest had a glass slide only.

In each trial, colonies were presented with two candidate nests, which differed in the number and types of attributes, as illustrated in Table 1. Six pairs of possible variable combinations were tested (n = 10) (Table 1). In Tests 1 and 2, we manipulated only one attribute to evaluate the effects of cavity *thickness* and entrance *width* in isolation. Tests 3 and 4 involved manipulating two nest attributes simultaneously. In Test 3, both *brightness* and *thickness* were manipulated to change in the same direction (i.e., in agreement). Nest 1 was dark and thick, both preferred attributes, while Nest 2 was bright and thin. In Test 4, both *thickness* and *width* were manipulated and in conflict. Additionally, Test 4 forced the ants to compare two lower-valued attributes by eliminating the attribute that was expected to be prioritized (brightness). In Tests 5 and 6, all three nest attributes were manipulated, and both manipulations were conducted such that there was no single ‘best’ nest option, but the nest attributes were always in conflict. In Test 5, Nest 1 was better than Nest 2 in *brightness* but less attractive in the other two attributes, with a thin cavity and a wide entrance. This allowed us to test whether the brightness outweighs the other two attributes when nests vary along all three. In Test 6, Nest 1 was better than Nest 2 in *brightness* and *width,* but had a thin cavity. This test acted as a more cognitively challenging version of Test 3, in which each nest choice had various attributes in conflict.

**Table 1.** Nest pairs tested. In all tests, Nest 1 was considered better than Nest 2 based on findings from previous studies. The differences between nests are bolded. All tests were done with 10 colonies.

| Test No. | No. of Differences | Nest 1 | Nest 2 |
| --- | --- | --- | --- |
| 1 | 1 | Dark, thin, & **narrow** | Dark, thin, & **wide** |
| 2 | 1 | Dark, **thick**, & wide | Dark, **thin**, & wide |
| 3 | 2 | **Dark, thick**, & wide | **Bright, thin**, & wide |
| 4 | 2 | Bright, **thick**, & **wide** | Bright, **thin**, & **narrow** |
| 5 | 3 | **Dark**, **thin**, & **wide** | **Bright**, **thick**, & **narrow** |
| 6 | 3 | **Dark**, **thin**, & **narrow** | **Bright**, **thick**, & **wide** |

### Experimental setup

The experimental arena consisted of two candidate nests, placed at an equal distance (approximately 7 cm) from the home nest, within a covered petri dish (Fig 2a). We placed a timer next to the nests to indicate the trial’s duration. To stimulate the initial emigration, we removed the cover from the home nest and placed an LED lamp 10 cm away to create a bright environment. We recorded emigration events using a GoPro camera, which filmed the trial in time-lapse (pictures taken at 2-s intervals) from a top-down angle (Fig 2b).

**Figure 2.**
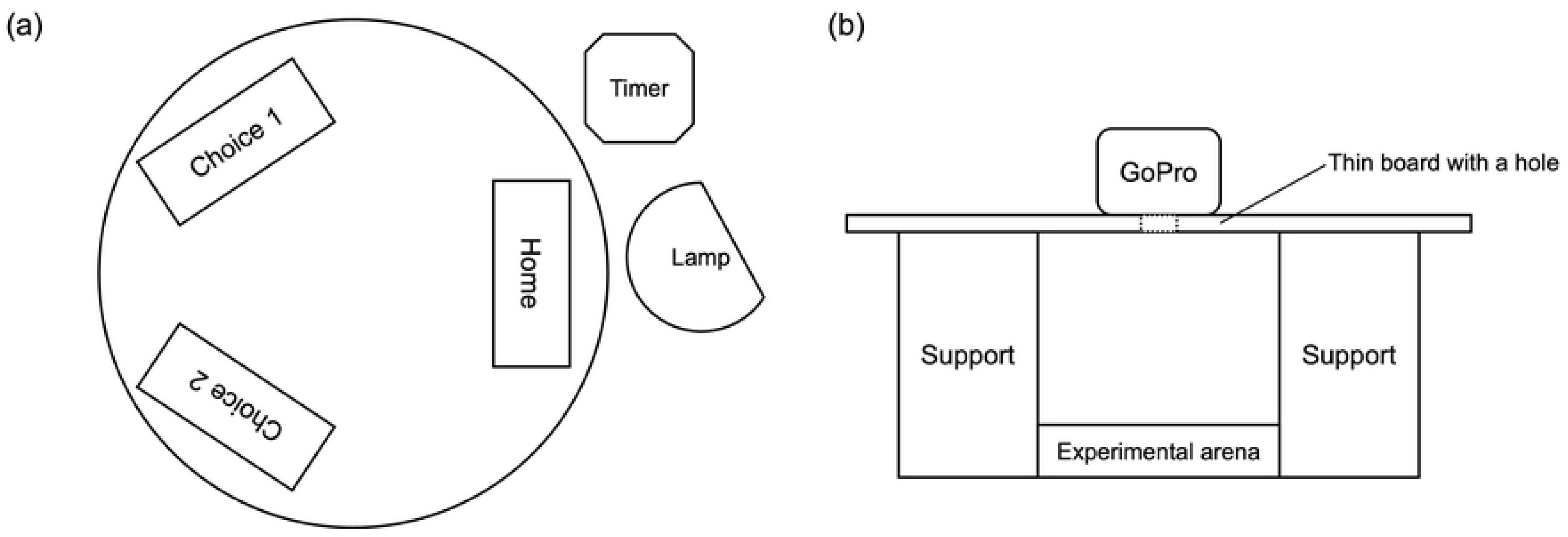
Set-ups of (a) experimental arena and (b) filming equipment (side view). Two candidate nests were placed approximately 7 cm from the home nest. The entrances of the nests all faced the center of the experimental arena. The lens of the GoPro faced down to film the arena from a top-down angle.

### Experimental protocol

Each colony (n = 10) encountered all six choice tests (Table 1). We randomized the order of presentation within and across colonies to minimize the chance that the ants would learn from their emigration experience (23,24). Prior to testing, colonies were housed in a home nest with the best possible attributes (i.e., dark, thick, and with a narrow entrance). To initiate a trial, the roof of the home nest was removed outside the arena to simulate a destroyed home nest and encourage emigration. A trial lasted until all the ants and brood were moved into a chosen nest, or the colony did not actively search for a new nest or initiate emigration after 1 hour. We recorded 1) the nest selected, 2) the duration between the first scout leaving the home nest and the beginning of direct transport (decision made), and 3) the duration between the first scout leaving the home nest and the end of the emigration event (decision finalized). While colony emigration is the point at which a decision is conclusively complete, we included both latency measures as colonies varied in size, and larger colonies might take more time to relocate all brood and individuals into the new nest, regardless of nest preferences.

Following a trial, each colony was placed back into the storage box with its restored home nest to allow the colony to return to its home nest before being retested. Colonies were each given at least 24 hours after returning to the home nest before being retested. It was important for this experiment that colonies always started from the same nest type, as ants may incorporate experience with previous nests when evaluating new nest sites (24).

### Data analysis

Analyses were carried out using R v. 4.3.1 (25). We used generalized linear models (GLMs) with the glm() and glmer() functions in the ‘lme4’ package (26). We first addressed whether nest preferences varied according to the number of attributes differentiating potential nest sites or whether preferences were variable between the test conditions, depending on the types of attributes manipulated. To test for the effects of number and type of attributes on nest preferences, we ran binomial GLMs (link = logit) with nest type (Nest 1 or Nest 2) as the response variable and either ‘attribute number’ or ‘test type’ as predictor variables. For our model testing the number of attributes, we did not include ‘colony’ as a random factor due to a singularity issue. To determine whether the number or type of attributes influenced either measure of decision latency, we ran GLMERs, now with a gamma distribution (link = log), commonly used for duration data with a positive right skew. We again included either ‘attribute number’ or ‘test type’ as the predictor variable, and ‘colony’ as a random factor. We used the Anova() function in the ‘car’ package (27) to evaluate model fit by generating Wald chi-square tests for categorical fixed effects. Where we found significant effects of a predictor variable, we used the function and package ‘emmeans’ for *post hoc* comparisons (Tukey’s honestly significant difference HSD tests) (28). A p-value of 0.05 is considered the threshold of significance.

## Results

### Choice accuracy

Increasing the number of attributes that differentiated potential nest sites significantly increased the probability that Nest 1 would be selected over Nest 2 (*χ*^2^ =10.089, df = 2, p = 0.006; Fig 3). Increasing from one to two attributes did not significantly increase choice accuracy for Nest 1 across the choice tests (z = - 1.671, p = 0.217), nor did increasing from two to three (z = −1.284, p = 0.404). However, nest accuracy significantly increased with three attributes relative to one attribute (z = −2.915, p = 0.01). The accuracy also differed significantly across the six test types (*χ*^2^ =24.135, df = 5, p < 0.001; Fig 3). These differences are most extreme between tests 2 and 3 (z = −2.574, p = 0.104) and tests 2 and 6 (z = −2.574, p = 0.104). In Test 2, these results were in the opposite direction from what was expected, with ants preferring the nest with the thinner cavity over the thicker. In Tests 3 and 6, preferences were almost exclusively for the brighter nest, regardless of other attribute values.

**Figure 3.**
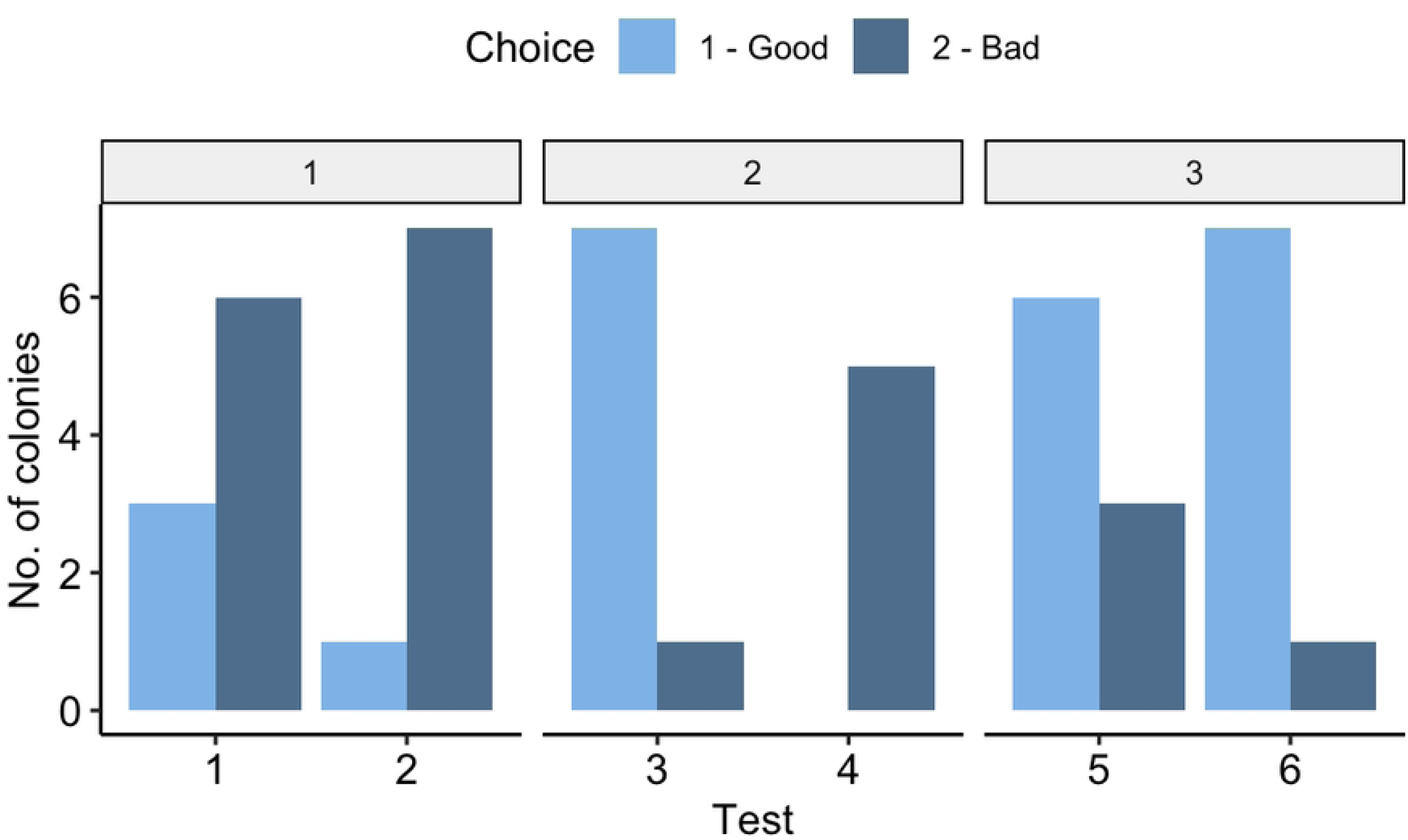
Choice accuracy across test types and the number of differences. Choices between potential nests depend on the number of differences between nest pairs and the six test types. The attribute number is denoted at the top of the figure (in the grey box), and the test type is plotted along the x-axis. Nest 1 was designed to be more attractive than Nest 2. The number of differences and the test type significantly affected which nest the colonies chose.

### Choice latency

Neither attribute number (χ^2^ = 3.695, df = 2, p = 0.158) nor test type (χ^2^ =3.709, df = 5, p = 0.592) affected the latency until direct transport (Fig 4). There was also no effect of attribute number (χ^2^ = 1.464, df = 2, p = 0.481) or test type ((χ^2^ = 2.111, df = 5, p = 0.834) on the latency until transport was complete.

**Figure 4.**
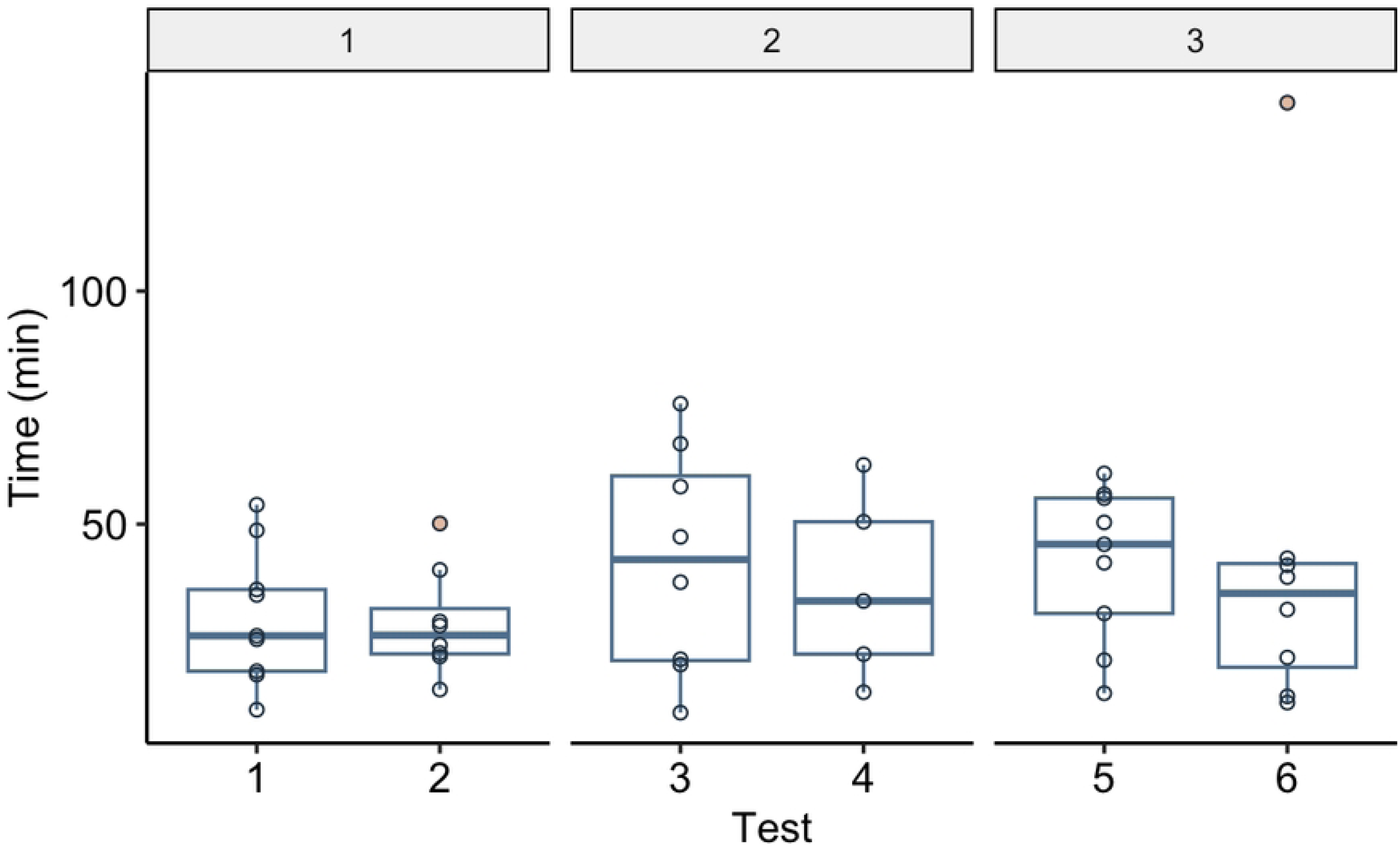
Latency until direct transport (decision made) across test types and the number of differences. Neither test type nor attribute number significantly affected the latency.

## Discussion

While multi-attribute decision problems are common across diverse taxa and decision-making contexts, the mechanisms by which animals make such complex decisions in many ecological settings remain unclear (4). Here, we explored how the number (one, two, or three) and type (brightness, width of entrance, and thickness) of attributes differentiating potential nest sites in acorn ants (*Temnothorax curvispinosu*s) influenced the *accuracy* and *latency* of nest-site selection. We found that the degree of difference between candidate nests affected the quality of the nest selected, but preferences for the more attractive nest were primarily explained by differences in brightness. We also found that the decision-making latency of colonies was similar across all choice contexts, regardless of attribute number or test type. Here, colonies spent, on average, around 68 minutes completing the emigration process, which is shorter than results found in previous experiments (7,12). These differences could possibly be explained by variation in the size of colonies or the distance between home nests and new nests. Together, our results suggest that ants may be temporally constrained when choosing a nest site, and the consequences of poor decisions may increase with the number and type of attributes differentiating potential nests.

Increasing the number of attributes differing between two potential nests led to more colonies selecting the preferred nest types. These results may be explained in several non-mutually exclusive ways. The first is that increasing the differences between potential nests may help facilitate decisions for the ‘better’ nest, even when some attributes conflict. Various drift-diffusion models address how animals make decisions when confronted with multiple options (e.g., (5)). One such model is called the ‘tug-of-war model’ in which an individual comparatively evaluates two options, computing an index of relative attractiveness between them, which is used to rank both options (20,29). The larger the difference between the two, the easier (and faster) a decision threshold is reached. Here, nests that varied in all three attributes may have facilitated this comparison, with more attributes creating greater contrast between two potential nests. Another explanation is that when nests vary slightly (i.e., only in one attribute), the fitness consequences of choosing an inferior option are small relative to the decision costs (regarding time or cognitive processing) (12,30). This may be especially true in the current study, as nests that varied in one attribute either varied in cavity thickness or entrance size, which are less preferred attributes (9). In either case, the fitness consequences of choosing the less-preferred nest are likely quite low.

Although attribute *number* influenced decisions between nest sites, attribute *type* appeared to be more important. When potential nests differed in only one attribute, cavity thickness or entrance width, ants did not select the nest we expected to be more attractive based on previous findings (9). These two attributes appear to be generally less prioritized, although previous studies have shown that ants still show robust preferences when these attributes are manipulated in isolation (9,14). Interestingly, preferences were detected in the opposite direction when thickness was manipulated in isolation: ants seemed to prefer the nest with the thinner cavity when thickness was the only attribute manipulated. These results may be explained by the possibility that we inadvertently reduced the internal nest space by manipulating cavity thickness and controlling for nest volume. In addition to the three attributes manipulated here, cavity volume may represent another attribute ants consider when selecting a potential nest site. Future studies could investigate whether and to what extent ant colonies consider this attribute when selecting potential nests.

Acorn ants are ecological engineers capable of transforming acorns and other nuts into suitable nesting habitats for their colonies. They can manipulate certain structural features of the nest, for instance, by excavating the interior to increase volume and modifying entrance holes to regulate access (31,32). They can even create an additional entrance to a cavity (32). Other aspects of potential nest sites, like internal brightness, however, are fixed and cannot be altered once a nest is chosen (9). As a result, acorn ants may place greater value on fixed attributes during nest selection, prioritizing them over attributes they can modify post-occupation. Across all choice comparisons, we found that ants prioritized nest brightness over the cavity thickness or the entrance’s width. Such findings agree with previous work, which suggests that brightness is the most prioritized attribute ants attend to when selecting a new nest site (9,12,14,20). In all test types where nest brightness was manipulated, colonies almost always chose the dark nest, regardless of other attribute differences. Although we didn’t have a manipulation where brightness varied independently, based on previous findings, we would expect ants to prefer the darker nest (9,12,14,20).

Unlike choice accuracy, decision speed was not affected by the number or type of attributes differentiating nests. Ant colonies chose a new nest within a similar period across all test conditions. These results contrast with those in a previous study exploring the temporal aspect of this decision task, which found that colonies made faster decisions when two nests were more similar (12). Given how variable preferences were across our test conditions, this result was a bit surprising, as we would have expected nests that differ in more attributes and, importantly, in brightness, would have facilitated faster decisions (29,30). Given that we only tested 10 colonies, we may have been limited in our ability to detect subtle differences in decision-making latencies between nests based on attributes; however, even if slight differences exist, rapid emigration appears critical, as ants are vulnerable to predators and larval desiccation without a nest. This suggests that ants prioritize making rapid decisions, even at the cost of potentially less accurate decisions. In such situations, selecting any potential nest site may offer greater protection than remaining exposed on the forest floor for an extended period.

This has interesting implications for the potential fitness consequences of nest selection in acorn ants. Variation in the accuracy of decisions, but not the time spent making them, suggests that there may be a speed-accuracy tradeoff in the nest-site selection process of ant colonies. If the task is complex and the cost of making decision errors is small, animals may choose randomly, resulting in lower choice accuracy (21). Notably, the critical factor in nest-site selection is not how different the nests are but rather what specific differences exist between them. Given the complexity of information processing required for decision-making, particularly in unpredictable environments, prioritizing rapid decisions over prolonged deliberation may be more advantageous for survival (34). Inhabiting a poor nest, which still provides temporary shelter and a chance to find a new one in the future, is better than having a broken nest where they are entirely exposed to the environment and perhaps predators.

Notably, only half of the colonies emigrated in the fourth test condition, all of which picked Nest 2, which was less ideal, being bright, thin, and narrow. It is possible that some colonies perceived Nest 2 as a better option because of the wider cavity space, as previously mentioned. Although the home nest had a thick cavity and narrow entrance, we removed the glass cover of the home nest to motivate emigration, completely exposing the internal cavity. Even though the new nests were also bright, they still had glass covers attached, and some colonies may have considered them better than their home nest, which was exposed entirely. Another non-mutually exclusive possibility is that ants experienced choice overload, a phenomenon in which decision quality deteriorates with an increasing number of options. This typically occurs when decision makers are confronted with more options, but it may also occur as options vary along multiple attributes (35). One of the symptoms of choice overload is choice deferral, where a decision is put off until a later time (35). In humans, one driver of this phenomenon is assortment complexity, where available options are similar and no dominant option exists (35,36). A previous study has explored choice overload in another context in house-hunting ants, and has shown how the number of potential nests influences decision-making in acorn ants, and found that individual ants experience choice overload (37). In their study, they found that when increasing the number of potential nests from two to eight, individual ants stopped selecting preferred nests and began choosing randomly (37). Future studies in ants and other animals could help determine how both the number of options and the attributes characterizing contribute to choice overload.

Because colonies arrive at a decision through a recruitment system that generates positive feedback, decisions are guided by interactions among ants rather than any single individual (8). Insect colonies show consistent preferences that are not simple summations of comparisons made by individual workers but instead emerge from interactions among individuals who only have knowledge of one available option (20,37,38). Previous studies have found notable differences between individual ants and colonies regarding the rationality of their decision-making. For instance, individuals are susceptible to choice overload and decoy effects, whereas colonies are not (19,37). Individual ants also take longer to complete an emigration when choosing between two similar nests than between two dissimilar nests, whereas colonies take less time when nests are similar (12). Finally, colonies outperformed individuals in a difficult perceptual task, but individuals performed better than groups when the task was easy (39). Such findings suggest that in our study, evaluating nest sites according to all three attributes and weighing those attributes accordingly may be a straightforward task at a colony level. However, individual ants performing the same task may show different patterns. Additionally, both individuals and colonies demonstrated repeatable variation in decision-making behaviors here, which could affect both the latency and accuracy of their decisions. Some colonies were more active, with scouts promptly searching for new nests and recruiting other ants, while other colonies were less active, only sending one or two scouts during the emigration process. In future studies, colonies and individuals could be categorized into different experimental groups based on their overall efficiency or activity to determine how variation at both levels influences decision-making behaviors.

Taken together, our findings suggest that the degree of differences does not affect the decision-making latency in house-hunting acorn ants, suggesting that ant colonies searching for a new nest might be constrained temporally when selecting a new nest site. However, differences influence the accuracy of decisions about which nests to emigrate to, particularly when nest brightness was manipulated, indicating that increasing the number of attributes facilitates nest-site selection. When multiple attributes characterize nests, this may have important consequences for decision accuracy and speed in ants. Such results highlight the importance of incorporating more complexity into choice studies to understand how animals make decisions in the wild.

## Acknowledgements

We would like to thank Dr. Collin Edwards and Dr. André Vieira Rodrigues for their help with data analysis, and Dr. Jessie Thuma for her input on the project idea.

## Supporting information

**S1 Table. Nest attributes in previous studies.** The table includes the dimensions of nests used in previous experiments. Pratt & Pierce (2001) used real acorns in the experiment. The values here are from the results after they measured the attributes of the acorns used.

**S1 Dataset. Dataset used in data analysis.** The data include all raw data collected in the experiment.

**S1 R Codes. R codes used in data analysis.** The R Notebook contains the codes used for data analysis.

